# Crossmodal metaperception: Visual and tactile confidence share a common scale

**DOI:** 10.1101/2021.07.07.451428

**Authors:** Lena Klever, Marie Mosebach, Katja Fiehler, Pascal Mamassian, Jutta Billino

## Abstract

Perceptual decisions are typically accompanied by a subjective sense of (un)certainty. There is robust evidence that observers have access to a reliable estimate of their own uncertainty and can judge the validity of their perceptual decisions. However, there is still a debate to what extent these meta-perceptual judgements underly a common mechanism that can monitor perceptual decisions across different sensory modalities. It has been suggested that perceptual confidence can be evaluated on an abstract scale that is not only task-independent but also modality-independent. We aimed to scrutinize these findings by measuring visual contrast and tactile vibration discrimination thresholds in a confidence forced-choice task. A total of 56 participants took part in our study. We determined thresholds for trials in which perceptual decisions were chosen as confident and for those that were declined as confident. Confidence comparisons were made between perceptual decisions either within the visual and tactile modality, respectively, or across both modalities. Furthermore, we assessed executive functions to explore a possible link between cognitive control and meta-perceptual capacities. We found that perceptual performance was a good predictor of confidence judgments and that the threshold modulation was similarly pronounced in both modalities. Most importantly, participants compared their perceptual confidence across visual and tactile decisions with the same precision as within the same modality. Cognitive control capacities were not related to meta-perceptual performance. In conclusion, our findings corroborate that perceptual uncertainty can be accessed on an abstract scale, allowing for confidence comparisons across sensory modalities.

## Introduction

We explore the world with multiple senses. What we perceive is the result of committing to perceptual decisions that are derived from uncertain sensory information. Along with these perceptual decisions usually comes a subjective, probabilistic estimate of how confident we are that this decision is correct (Fleming et al., 2012) or at least self-consistent (Mamassian, 2020). Typically, subjective confidence judgments and objective task performance are correlated (Fleming & Lau, 2014), but they can also diverge (Lau & Passingham, 2006). As confidence judgements are a second-order judgement on a first-order judgement, they are commonly summarized under the term “metacognition”. Metacognition can operate over a variety of domains – such as perception and cognition. It has been studied for decades but mainly in the domain of memory (Koriat, 2007). Only recently, newer, more sophisticated methods have been developed to investigate confidence judgments underlying perceptual decisions (Fleming & Lau, 2014; Mamassian, 2020). When it comes to the perceptual domain, the sub-categories “meta-perception” and “confidence” are used to specifically describe an observer’s ability to monitor, evaluate and control their own perception (Mamassian, 2016).

In everyday life, meta-perception is essential for guiding behavior (Desender et al., 2018). Especially in noisy environmental situations, such as fog, it might be helpful to rely on cues from different sensory modalities. Considering our confidence in these perceptual decisions might further be useful to adapt our behavior. For instance, if we’re not sure that a car is approaching, we would rather look twice before crossing the road. In addition, we could also reflect whether we heard an engine or felt the ground vibrating. Confidence from multiple senses seems to be helpful to select the sensory information that appears most reliable. However, this selection process would only be functional and economically useful if confidence could be easily compared across different modalities.

So far, confidence across the senses remains largely understudied. It is still an open question whether post-decisional confidence is not only task-independent but also modality-independent and can serve as a common currency across different modalities. A common currency, or a task-general view, suggests that meta-perceptual judgements for different tasks are stored in the same format (deGardelle & Mamassian, 2014). Due to this common currency, confidence comparisons across different tasks should be facilitated and be possible without any information loss. A task-specific view, in turn, suggests that there are different metaperceptual formats for different perceptual tasks (deGardelle & Mamassian, 2014). Consequently, these formats need to be transformed to allow for comparisons across tasks, resulting in conversion costs (e.g. information loss, time increase) and make the process more effortful.

Behavioral studies typically addressed the question whether confidence can be generalized across different tasks and modalities using a correlational approach (e.g. Faivre et al., 2018, Mazancieux et al., 2020, Morales et al., 2018). Here, the idea is that meta-perceptual performance in one task should be associated with meta-perceptual performance in another task if confidence was a general resource (in analogy to the g-factor for intelligence). Only few studies have tested this idea directly by asking participants to compare their confidence in one task with their confidence in another task (e.g. Baer & Odic, 2020, deGardelle et al., 2016). Despite these different approaches, findings from behavioral studies are mostly consistent and support task- but also modality-general mechanisms. For instance, it has been shown that observers can directly compare their confidence across two different visual tasks (deGardelle & Mamassian, 2014) and across a visual and an auditive task (deGardelle et al., 2016) without additional costs. A common confidence format even seems to be present in 6-7-year-old children for different perceptual tasks in which they had to judge emotional expressions or the size of a stimulus (Baer & Odic, 2020). But there have also been reports that challenge the idea of a common scale: Metaperceptual performance was only weakly correlated between a luminance and auditory discrimination task and did not correlate between a contrast and auditory discrimination task (Ais et al. 2016).

Though the first confidence study focused on the tactile sense (Pierce & Jastrow, 1884), the tactile modality has received less attention afterwards – especially in multisensory situations. Most studies examining crossmodal meta-perception focused on vision and audition (e.g. Ais et al., 2016, deGardelle et al., 2016). However, a shared common currency between the visual and the tactile sense appears particularly useful: In contrast to vision and audition, vision and touch often interact and are closely tied to one another when performing actions. This strong action-perception coupling might profit further from confidence as a common currency since actions need to be permanently adjusted to properly respond to newer incoming sensory information. Confidence about the quality of this incoming sensory information appears helpful to determine the reliability of sensory cues and guide further selection processes. If confidence served as a common currency between the visual and tactile sense, selection processes could even be sped up. So far, there’s some evidence for a common currency between the visual and tactile sense. For instance, Faivre et al. (2018) found that meta-perceptual ability in a vibrotactile discrimination task was associated with meta-perceptual ability in a contrast discrimination task. However, the two perceptual tasks were performed in separation and crossmodal interactions couldn’t be considered. It might be possible that interactions between the visual and tactile sense are influenced by the belief that touch provides more directness and certainty (Deroy & Fairhurst, 2019). Indeed, there is first evidence that observers favor touch over vision and report higher confidence in touch when the situation is ambiguous (Fairhurst et al., 2018). Such an overreliance on the tactile sense could challenge the idea of confidence as a common currency between vision and touch.

While findings from behavioral studies can provide first insights into whether meta-perception relies on a common mechanism, neuroimaging studies can examine the formation process of meta-perception in greater depth. More specifically, they can provide information on whether meta-perceptual processes for different sensory modalities involve similar brain regions and networks or brain regions that are specific for each modality. They have been proven useful to better understand conflicting behavioral findings: It appears that the two perspectives on meta-perception are not necessarily exclusive. There is evidence that both domain-specific and domain-general signals co-exist in the human brain, but their activation is task-dependent (Morales et al., 2018, Rouault et al., 2018). The most critical neural functional correlate of metacognition being involved irrespective of the first-order task is the (anterior) prefrontal cortex (Fleming & Dolan, 2012, Valk et al., 2016). In consistency with the involvement of the prefrontal cortex, metacognition and executive functions (EF) have been described as closely related constructs (Fernandez-Duque et al., 2000, Roebers, 2017). EF refers to effortful cognitive control operations, such as inhibition of irrelevant information, task-switching and updating (Miyake & Friedman, 2012). Both EF and metacognition are higher-order cognitive processes that mainly comprise monitoring and control abilities. So far, only few studies have considered links between EF and metacognition. The results are heterogeneous and primarily stem from aging research (e.g. Palmer et al., 2014).

Here, we investigated whether confidence serves as a common currency between the visual and the tactile sense - two senses that closely interact and are especially relevant for the planning and execution of actions. Given this close interaction, we wanted to directly examine how well observers can compare their confidence across a visual and a tactile task. To this end, we applied the well-established confidence forced-choice paradigm (deGardelle & Mamassian, 2014), where participants performed two perceptual tasks in succession and then selected the perceptual decision that they think is more likely to be correct. If confidence is modality-specific, we would expect that comparisons across perceptual modalities are harder than within the same modality. Conversely, if confidence was stored in a modality-general format, comparisons across modalities should be as precise as comparisons within the same modality. To evaluate the effortfulness of this meta-perceptual process, we further explored possible links to cognitive control capacities. Positive correlations between crossmodal meta-perceptual ability and cognitive control capacities might provide first insights into whether crossmodal meta-perceptual processes also draw on cognitive control resources.

## Methods

### Participants

A total of 56 participants (13 males) with a mean age of 24.1 years (*SD* = 5.8 years) took part in this study. All participants had normal or corrected-to-normal vision and no history of ophthalmologic, neurologic, or psychiatric disorders.

In addition, we used a battery of standard cognitive tasks to evaluate key facets of executive functions (EF) for each participant: Updating ability (Digit Symbol Substitution Test; Wechsler, 2008), shifting ability (Trail Making Test part B; Kortte et al., 2002, Reitan & Wolfson, 1985), inhibition ability (Victoria Stroop Test color naming; Mueller & Piper, 2014, Stroop, 1935) and nonverbal reasoning ability (LPS-2; Kreuzpointner et al., 2013). To obtain individual global EF scores, all measures were z-standardized and then averaged for each participant. As the procedure of our meta-perceptual task was rather complex, we additionally assessed the maximal backward digit span (Härting et al., 2000) to evaluate short-term memory capacity.

Methods and procedures were approved by the local ethics committee of the Faculty of Psychology and Sports Science, Justus Liebig University Giessen, and were carried out in accordance with the guidelines of the Declaration of Helsinki (World Medical Association, 2013). Participants were compensated with course credits or money.

### Setup

Visual stimuli were presented on a calibrated 32” Display++ LCD monitor (Cambridge Research Systems, Rochester, UK) with a spatial resolution of 1920 × 1080 pixels and a refresh rate of 120 Hz (non-interlaced) using the Psychophysics Toolbox (Brainard, 1997; Kleiner, 2010) in MATLAB (The Mathworks, Inc., Natick, MA, USA). The background was average grey. Participants sat at a table in a darkened room with their head stabilized on a chin rest. The eye-monitor distance was 100 cm, leading to a display size of 41° × 23°. Luminance of white and black pixels was 112.7 and 0.1 cd/m2, respectively, as measured with a CS-2000 Spectroradiometer (Konica Minolta). Tactile stimuli were applied by custom-made vibrotactile devices (Engineering Acoustics Inc., Casselberry, FL, USA). They were attached on the tip of both index fingers using silicone finger sleeves. Participants comfortably rested their hands shoulder-width apart on foam pads in front of them. Due to the setup for tactile stimulation, manual response input was excluded. Thus, we used gaze positions as response input. Eye positions were recorded using an SR Research Eyelink 1000 Desktop Mount system (SR Research Ltd., Mississauga, Ontario, Canada).

### Stimuli and procedure

Meta-perceptual ability was assessed in an established confidence forced-choice paradigm (Mamassian, 2020). An advantage of this approach is that it focuses on metaperceptual sensitivity, i.e. an observer’s ability to discriminate between correct and incorrect perceptual decisions, while minimizing confidence biases (Mamassian, 2020). One trial comprised two consecutive perceptual tasks (visual and/or tactile) and a confidence task (see Figure 1A). All possible order combinations of the two perceptual tasks were realized in four separate blocks with 112 trials each and considered the type of confidence comparison: unimodal (visual – visual, tactile – tactile) and crossmodal (visual – tactile, tactile – visual). Every participant completed all 4 blocks; block order was counterbalanced across participants. Prior to each block, participants completed 14 training trials of the complete procedure. After each block, they had the opportunity to take a break. Before data collection, we provided an extensive introduction to our procedure. In order to familiarize with the tasks and in particular with the response input by gaze, participants practiced the visual and tactile task, respectively.

**Figure 1.**
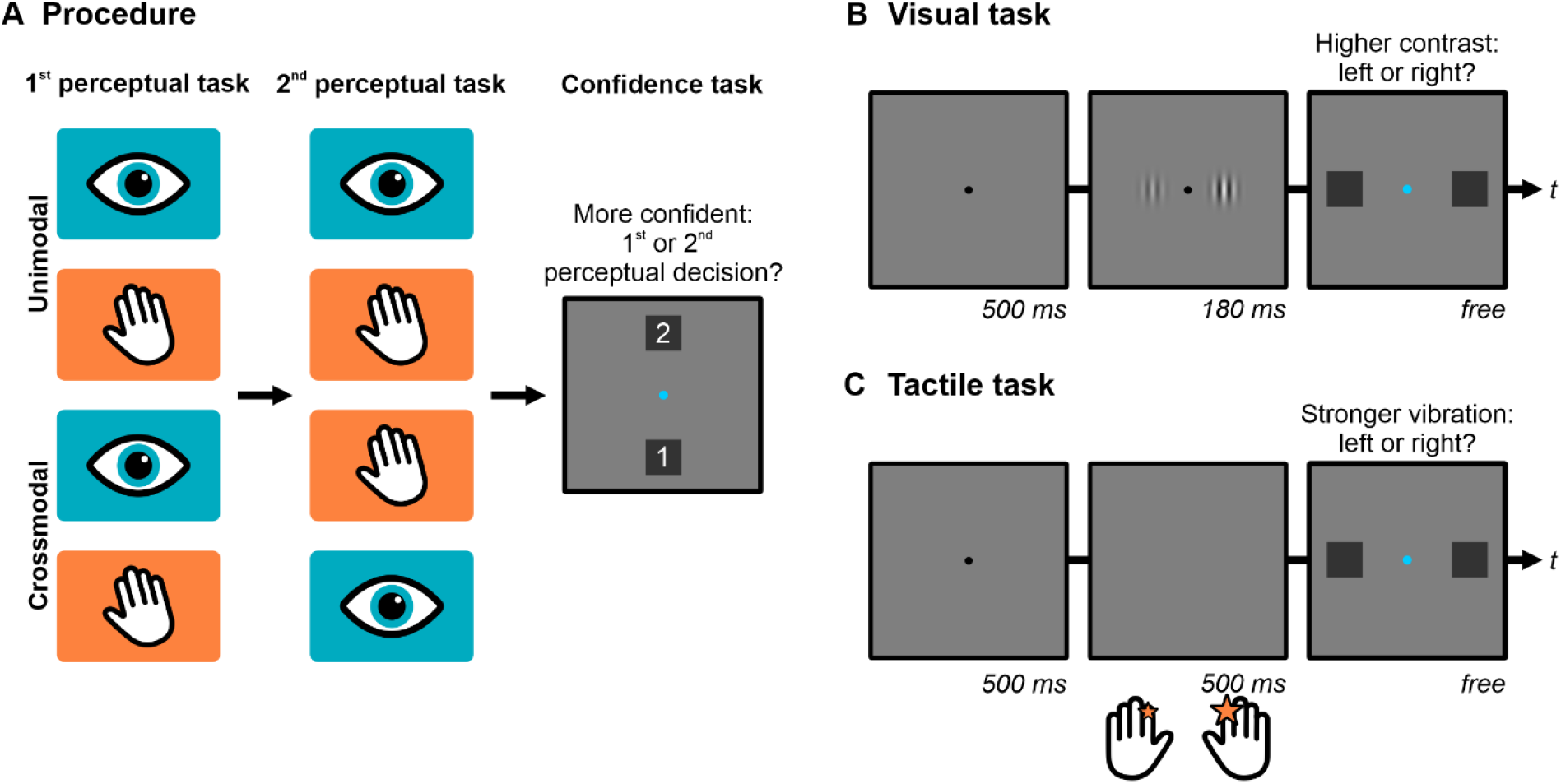
Procedure and subtasks of the confidence forced-choice paradigm. (A) Schematic illustration of the overall trial procedure. Participants completed two perceptual tasks in succession (either visual-visual, tactile-tactile, visual-tactile or tactile-visual) and then provided a forced-choice confidence judgement, i.e. they indicated about which of the two perceptual decisions (first or second) they felt more confident. (B) Visual task. Participants first saw a fixation dot, which was followed by the simultaneous presentation of two Gabor patches. Then, they decided which of the two Gabor patches appeared higher in contrast. (C) Tactile task. First, participants were presented with a fixation dot. Next, they received two simultaneous vibrations on both index fingers and decided afterwards on which finger the vibration felt stronger.

The visual task (Figure 1B) started with a 500 ms presentation of a central black fixation dot that subtended 0.2°. Then, two vertical Gabor patches were simultaneously shown for 180 ms on the left and right of the fixation dot at 4.2° eccentricity. All Gabor patches had a spatial frequency of 0.8 cyc/°. The standard deviation of the Gaussian envelope was 1° and the phase was randomized. Of these two Gabor patches, one always had a fixed contrast at 22% (standard Gabor patch), while the contrast of the other Gabor patch was adapted throughout the experiment (test Gabor patch). Laterality of standard and test Gabor patch was randomized. Next, the fixation dot turned blue and two dark-grey response squares were shown at 6.8° eccentricity left and right from the fixation dot. Participants’ task was to decide whether the left or right Gabor patch appeared higher in contrast by looking at the respective response square. When a square was selected, it turned darker. Based on participants’ decision, the contrast for the test Gabor patch of the next trial was adapted by one of two randomly interleaved 3-down/1-up staircases in steps of 3%: One staircase had a starting value of 31% and aimed at responses favoring the test stimulus ~80% of the time; the other had a starting value of 13% and aimed at responses favoring the standard stimulus ~80% of the time.

The tactile task (Figure 1C) began with the same fixation dot configuration as the visual task. Following, participants received two simultaneous vibrations for 500 ms on both index fingers at a frequency of 200 Hz. Of these two vibrations, one had a fixed intensity, defined as peak-to-peak displacement, of 0.13 mm (standard vibration). The intensity of the other vibration (test vibration) was adapted throughout the experiment. Again, laterality of standard and test stimuli was randomized. When the horizontal response squares were shown, participants had to decide whether the vibration on the left or right index finger felt stronger by looking at the according square. Using similar staircases to the visual task with starting values of 0.08 mm and 0.18 mm, respectively, the intensity for the test vibration of the next trial was adapted in steps of 0.02 mm.

After the completion of two perceptual tasks, a blue central fixation dot and two dark-grey response squares were shown at 6.8° eccentricity below and above the fixation dot. The response squares were numbered and associated with the first or second perceptual decision. The mapping was visualized on the screen and balanced across participants. By looking at one of the two squares, participants indicated about which perceptual decision (first or second) they felt more confident (confidence judgement).

### Data analyses

For all combinations of modality (visual/ tactile) and comparison type (unimodal/ crossmodal) and each participant, we extracted all trials that were chosen in the confidence task as relatively more confident. Hence, these sets of trials are labeled as *chosen*. A second set of trials included all perceptual decisions irrespective of participants’ confidence for each combination. These sets are therefore labeled as *unsorted*.

Perceptual performance was then evaluated separately for each confidence set and condition by fitting cumulative Gaussian functions to the percentage of responses in which observers favored the test stimulus over the standard stimulus. The inverse standard deviation of the fitted psychometric functions provides a measure of sensitivity. We used the Psignifit 4 toolbox in Matlab for the fitting process, as it yields an accurate estimation of psychometric functions in a Bayesian framework even if the measured data is overdispersed (Schütt et al., 2016). Goodness of fit of the psychometric functions was assessed with the measure of deviance *D*, which supported good fits between the model and the data. By inspecting boxplots for the derived sensitivity measures, we identified two participants who showed visual or tactile sensitivities that deviated more than 1.5 times the interquartile range from the range borders. We considered these measures as outlier data and discarded the participants from further analyses in order to reduce unsystematic noise in our data.

To analyze meta-perceptual sensitivity, i.e. the ability to estimate the accuracy of a perceptual decision, we calculated a confidence modulation index (CMI) according to Equation 1. The CMI quantifies meta-perceptual ability as the gain in sensitivity from the set of unsorted trials to the set of chosen trials standardized by the sensitivity derived from the unsorted trials. Thus, CMIs will increase with better metaperceptual sensitivity. If an individual observer shows low metaperceptual sensitivity, CMIs will be close to zero. Importantly, as a proportional measure, the CMI allows to compare meta-perceptual sensitivity across both modalities. CMIs were arcsine-square-root transformed for variance stabilization.

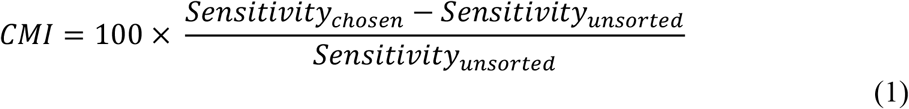

Time measures for perceptual decisions were explored using median response times (RT). For all analyses, only RTs between 150 ms and 3000 ms were considered in order to reduce noise from outlying values. Perceptual decision times might be confounded with stimulus difficulty – i.e. easier stimuli lead to faster response times and are more likely to be chosen as confident. To separate effects of stimulus difficulty and confidence on response times, we applied an elaborate analysis that was successfully used in previous studies (e.g. deGardelle & Mamassian, 2014, deGardelle et al., 2016). First, we normalized stimulus values for each participant, confidence set, modality and comparison type. This was realized by calculating the signed distances S between the stimulus intensities and the respective point of subjective equality in standard deviation units of the psychometric function. Next, we divided the normalized stimulus values into 5 bins and calculated the median response time as well as the average confidence judgement C (encoded as 0 for unchosen and 1 for chosen perceptual decisions) for each bin. Then, we fitted an exponential model with three free parameters (as defined by Equation 2) to the median RTs, separately for each condition. The estimated parameters are the following: α provides the generic RT, β reflects the exponential change in RT due to stimulus difficulty, and γ captures the linear change in RT due to confidence.

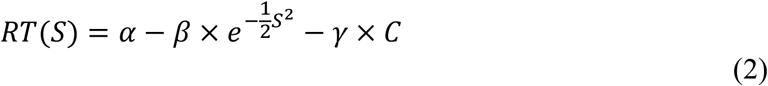

Perceptual sensitivities and RT were analyzed separately for each modality using repeated measures ANOVAs with the within-subject factor *confidence set* (chosen vs. unsorted) and the within-subject factor *comparison* (unimodal vs. crossmodal). To compare meta-perceptual sensitivity across modalities, we submitted CMIs to a repeated measures ANOVA with the within-subject factor *modality* (visual vs. tactile) and *comparison* (unimodal vs. crossmodal). Two-sided *t*-tests were used to further analyze CMIs and RT parameters. In case of unequal variances as indicated by Levene’s test, degrees of freedom were adjusted. Associations between metaperceptual sensitivity and cognitive measures were scrutinized by linear regression analyses. For all statistical analyses, a significance level of α = .05 was applied. Descriptive values are reported as means ± 1 SEM, unless stated otherwise.

## Results

First, we explored response patterns across all combinations of modality (visual vs. tactile) and comparison type (unimodal vs. crossmodal). Response patterns might provide first insights into potential biases and meta-perceptual performance. Then, we evaluated the effects of modality and comparison type on perceptual and meta-perceptual performance in detail by analyzing perceptual sensitivity functions for the chosen and unsorted confidence sets. Finally, we analyzed differences in response times and explored individual differences in cognitive resources that might contribute to meta-perceptual performance.

### Response patterns

The design of the confidence forced-choice paradigm ensured that the number of as confident chosen trials is equal in the unimodal conditions (112 trials). For the crossmodal comparisons, the number of chosen trials can vary due to biases towards one modality. On average, there were 105.13 ± 3.29 chosen trials in the visual crossmodal condition and 118.87 ± 3.29 chosen trials in the tactile crossmodal conditions. However, this slight bias towards the tactile modality should not have impaired the estimation of the psychometric functions as the measure of deviance supported good fits between the model and the data.

A crucial aspect of meta-perceptual performance is the degree to which an observer can distinguish correct trials from incorrect trials. Typically, participants will report higher confidence when their perceptual decision was objectively correct and lower confidence when their decision was objectively incorrect. Figure 2 illustrates average confidence judgments for correct and incorrect perceptual decisions at different normalized stimulus intensity levels.

**Figure 2.**
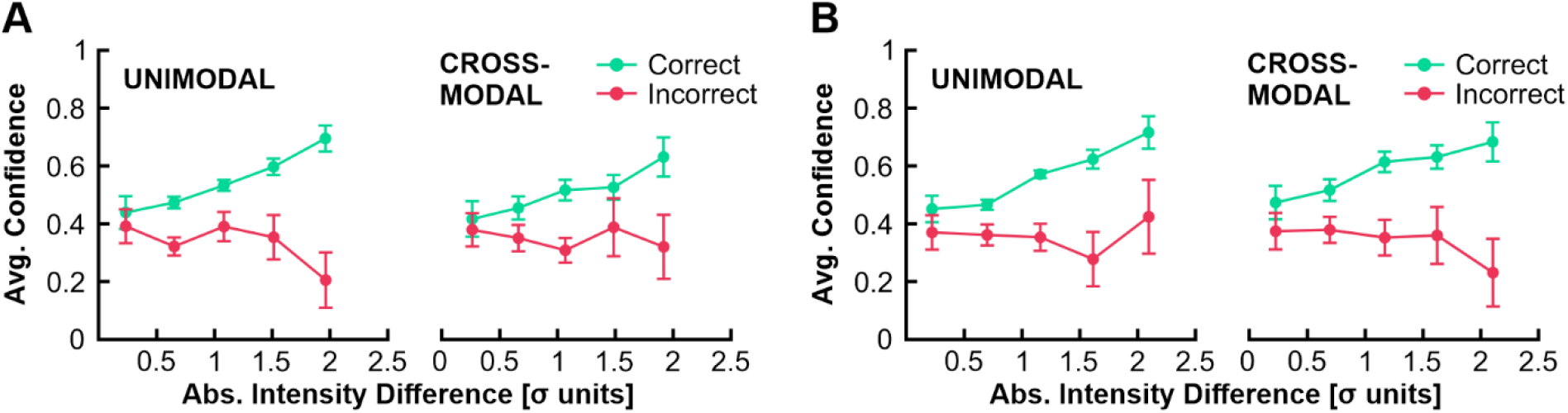
Average confidence judgements for correct (green) and incorrect (red) perceptual decisions at different intensity levels in the visual task (A) and tactile task (B), separately for the confidence comparison type (unimodal vs. crossmodal). Intensity levels are given as the absolute difference between stimulus intensities and each participant’s point of subjective equality in standard deviation units of the psychometric function. They were then divided into five bins of varying stimulus difficulty with higher values indicating lower stimulus difficulty. Confidence judgements were coded as 1 for chosen and 0 for unchosen perceptual decisions. Error bars provide 95% confidence intervals.

The overall response pattern suggests that participants evaluated their perceptual performance appropriately, irrespective of condition: Average confidence judgments were consistently higher for correct than incorrect trials. Additionally, the difference in average confidence judgements between correct and incorrect trials became more evident with decreasing stimulus difficulty.

### Perceptual performance

We were interested in determining whether sensitivities vary systematically between the chosen and unsorted trial sets, as well as unimodal and crossmodal comparisons. As sensitivities cannot be compared across modalities, we submitted sensitivity data separately for each modality to a repeated measures ANOVA with the within-subject factors *confidence set* (unsorted vs. chosen) and *comparison type* (unimodal vs. crossmodal). This analysis can provide further insights into meta-perceptual performance: If participants show some meta-perceptual ability, they should be able to select the trial that led to a higher performance. This, in turn, would result in higher sensitivities for the chosen trial sets. Indeed, the analysis yielded a strong main effect of *confidence set* for the visual task, *F*(1,53) = 109.42, *p* < .001, η_p_^2^ = .67, as well as the tactile task, *F*(1,53) = 172.25, *p* < .001, η_p_^2^ = .77. Sensitivities were consistently higher for the chosen confidence set in comparison to the unsorted confidence set. Figure 3 shows representative psychometric functions for contrast discrimination (A) and vibrotactile intensity discrimination (B). Of special interest is the observation that the psychometric functions derived from the chosen and unsorted trial set differ in slope, while the points of subjective equality are similar.

**Figure 3.**
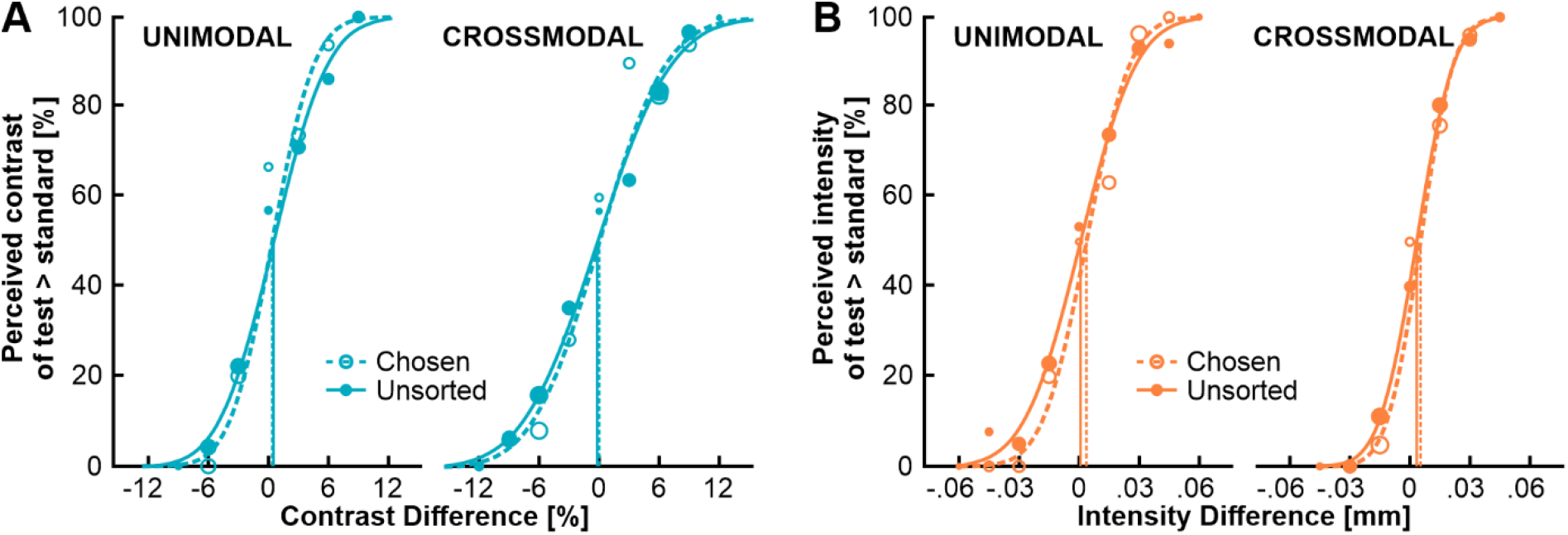
Representative psychometric functions of contrast discrimination (A) and vibrotactile intensity discrimination (B) for both unimodal and crossmodal comparison types. The proportion of choosing the test stimulus over the standard stimulus is plotted as a function of stimulus intensity given as the difference between the test and standard stimulus. Dashed lines and open dots depict data from the as confident chosen trial set, solid lines and filled dots represent data from the unsorted trial set.

Furthermore, there was a main effect of *comparison type* for the visual task, *F*(1,53) = 4.65, *p* = .036, η_p_^2^ = .08, but not for the tactile task, *F*(1,53) = 0.03, *p* = .857, η_p_^2^ < .01. Visual sensitivity was overall higher when derived from unimodal trials. This indicates that unimodal comparisons were slightly easier than crossmodal comparisons in the visual task. The interaction between *confidence set* and *comparison type* did not reach significance in either modality, visual: *F*(1,53) = 0.01, *p* = .947, η_p_^2^ < .01, tactile: *F*(1,53) = 1.20, *p* = .394, η_p_^2^ = .02, suggesting that meta-perceptual ability was not affected by the type of confidence comparison. Figure 4 illustrates the effects of confidence set and comparison type on sensitivities separately for the visual task (A) and tactile task (B).

**Figure 4.**
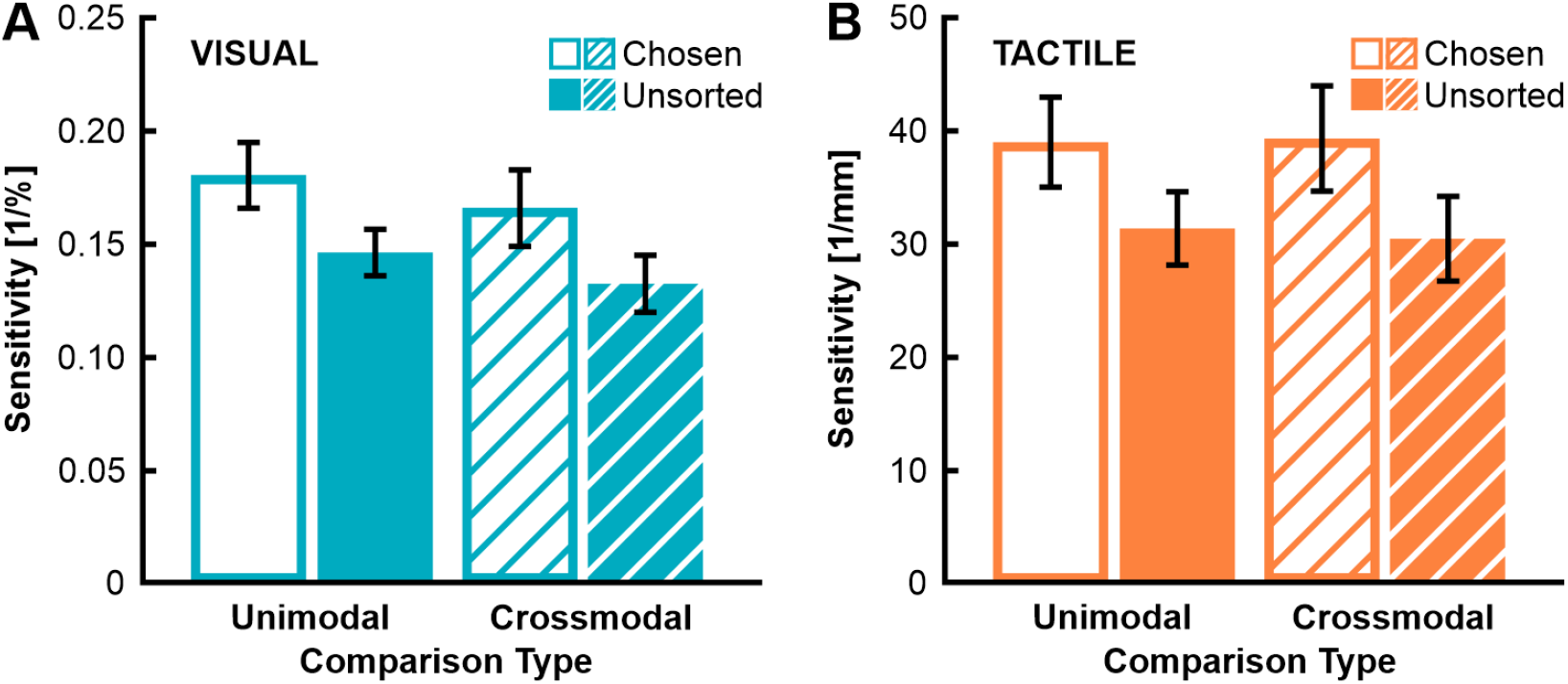
Average perceptual sensitivity as a function of confidence set and comparison type, separately for the visual task (A) and tactile task (B). Open bars represent mean sensitivities from the as confident chosen trial set, filled bars show mean sensitivities from the unsorted trial set. Hatched bars represent the crossmodal conditions. Error bars indicate 95% confidence intervals.

### Meta-perceptual performance

As the measures of sensitivity do not allow a direct comparison between the visual and tactile task, we further analyzed effects of modality and comparison type on meta-perceptual performance with the help of a Confidence Modulation Index (CMI; see Methods). For the visual task, the average CMI was 26.03 ± 2.11 in the unimodal condition and 28.90 ± 2.50 in the crossmodal condition. For the tactile task, the average CMI was 28.96 ± 1.57 in the unimodal condition and 31.90 ± 2.57 in the crossmodal condition. CMIs were consistently greater than zero in all conditions (all *p*’s < .001). Figure 5 displays average CMIs for both modalities and comparison types.

**Figure 5.**
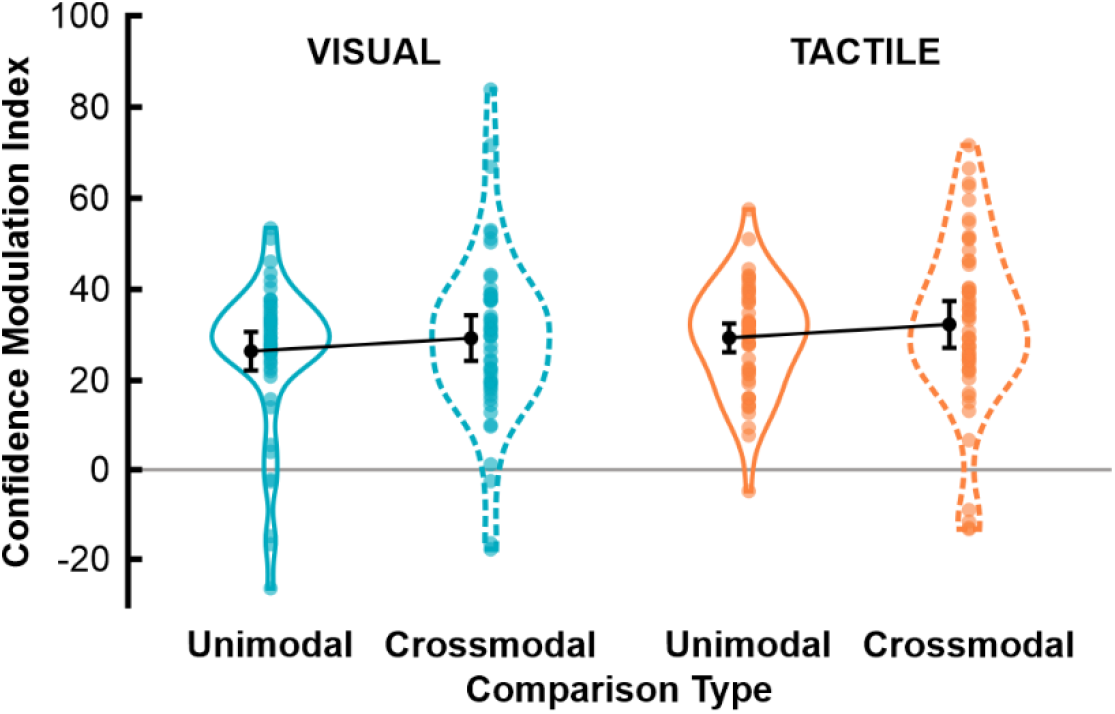
Confidence Modulation Index (CMI) as a function of modality (visual in blue, tactile in orange) and comparison type. The CMI is a proportional measure reflecting the change in sensitivity from the set of unsorted trials to the chosen trials relative to the unsorted trials. The higher the CMI, the higher the meta-perceptual ability. Colored dots represent individual data points; black dots display the mean across observers with error bars indicating 95% confidence intervals. Solid (unimodal) and dashed (crossmodal) outlines show 95% of the data distribution smoothed by a kernel density function.

We submitted CMIs to a repeated measures ANOVA with the within-subject factors *modality* (visual vs. tactile) and *comparison type* (unimodal vs. crossmodal). The main effect of *modality* was not significant, *F*(1,53) = 2.28, *p* = .137, η_p_^2^ = .04, indicating comparable meta-perceptual performance in both modalities. In line with the assumption that confidence is stored in a task-independent format, there was no main effect of *comparison type*, *F*(1,53) = 2.50, *p* = .120, η_p_^2^ = .05. The interaction between *modality* and *comparison type* was not significant, either, *F*(1,53) < 0.01, *p* = .986, η_p_^2^ < .01. Bayesian statistics provided support for the null hypothesis. There is evidence that neither *modality*, *BF*_10_ = 0.42, nor *comparison*, *BF*_10_ = 0.41, in isolation, nor their interaction, *BF*_10_ = 0.19, have an effect on meta-perceptual performance.

### Response times

Finally, we analyzed whether response times (RT) vary systematically between the confident chosen and unsorted trial sets and whether the type of comparison influenced RTs. To this end, we conducted a repeated measures ANOVA on median RTs separately for both perceptual tasks with the within-subject factors *confidence set* (unsorted vs. chosen) and *comparison type* (unimodal vs. crossmodal). We found a main effect of *confidence set* in the visual task, *F*(1,53) = 29.87, *p* < .001, η_p_^2^ = .36, as well as the tactile task, *F*(1,53) = 43.01, *p* < .001, η_p_^2^ = .45, indicating faster responses with higher confidence. Additionally, there was a main effect of *comparison type* in both modalities; visual: *F*(1,53) = 74.79, *p* < .001, η_p_^2^ = .59, tactile: *F*(1,53) = 6.71, *p* = .012, η_p_^2^ = .11. However, the direction of the effect differed between both modalities: In the visual task, responses were faster in the unimodal condition relative to the crossmodal condition. Whereas in the tactile task, responses were slightly faster in the crossmodal condition compared to the unimodal condition. Responses were made via saccades on a screen, which might have caused this asymmetry. There was no interaction between *confidence set* and *comparison type* in the visual task, *F*(1,53) = 0.09, *p* = .768, η_p_^2^ < .01, or tactile task, *F*(1,53) = 0.23, *p* = .633, η_p_^2^ < .01. Figure 6 illustrates effects of confidence set and comparison type on median response times separately for each modality.

**Figure 6.**
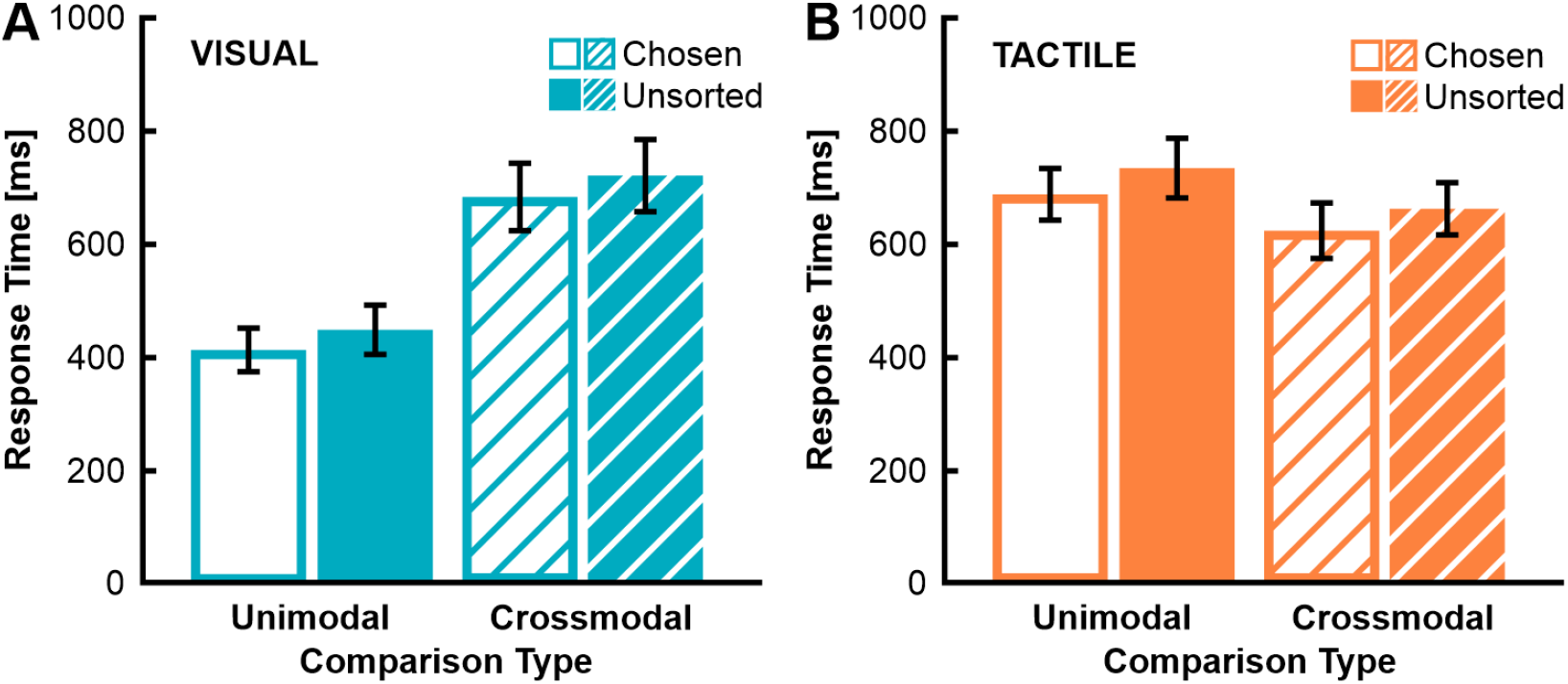
Median response times as a function of confidence set (chosen vs. unsorted) and comparison type (unimodal vs. crossmodal) for the visual task (A) and tactile task (B). Bars show the mean of observers in each condition with open bars representing the chosen trial set and filled bars the unsorted trial set. Hatched bars represent the crossmodal conditions. Error bars indicate 95% confidence intervals.

However, as these results might be confounded with stimulus difficulty, we modeled the relationship between stimulus difficulty, confidence level and response times (see Methods for details and Fig. 7). This control analysis yielded three parameters, but only α (the generic RT) and γ (the confidence effect) are relevant for the comparison with the previous analysis. The control analysis supported the finding that responses were slower in the visual crossmodal condition compared to the visual unimodal condition, *t*(53) = 2.45, *p* = .017. Interestingly, there was also a trend for slower crossmodal RTs than unimodal RTs in the tactile task, *t*(53) = 1.88, *p* = .066. In contrast to the previous analysis, the confidence effect was only greater than zero in the visual-crossmodal condition, *t*(53) = 2.30, *p* = .25. Thus, only in the visual-crossmodal condition, responses for the chosen trials were faster than the unsorted trials, when taking the effect of stimulus difficulty into account. For all other conditions, γ did not differ significantly from zero (all *p*’s > .289).

**Figure 7.**
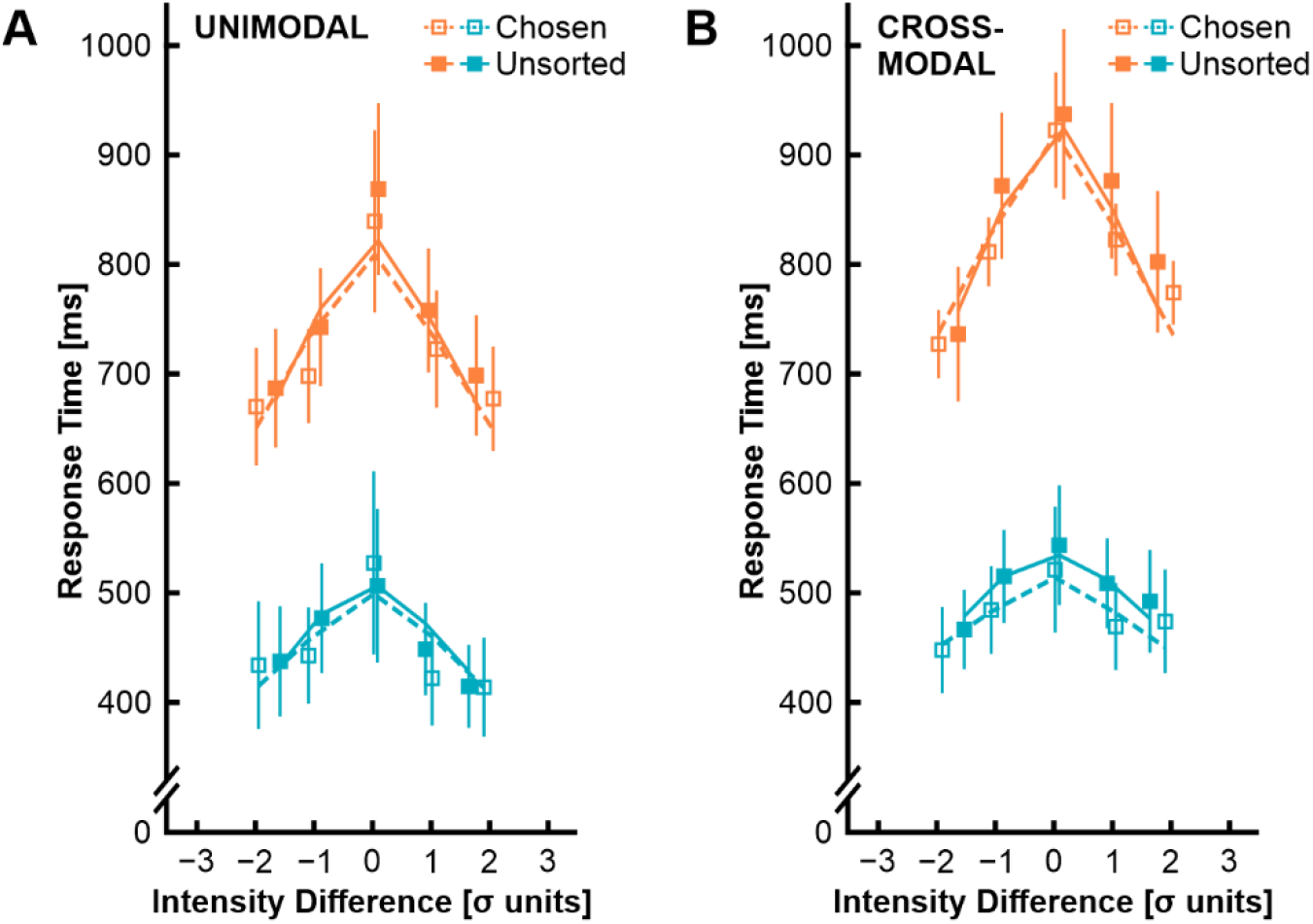
Additional analysis controlling for the effect of stimulus difficulty on response times. Median response times for five bins of stimulus intensities (in standard deviation units of the psychometric function) separately for the unimodal (A) and crossmodal (B) condition. Data from the tactile task is shown in orange, the visual task in blue. Open squares (chosen trial set) and filled squares (unsorted trial set) depict mean response times across observers with error bars showing 95% confidence intervals. Dashed lines (chosen trial set) and solid lines (unsorted trial set) show the average response time in each bin as predicted by our RT model.

### Individual differences in meta-perceptual sensitivity

Since we observed substantial variability in meta-perceptual sensitivity, especially in the crossmodal conditions (see Fig. 5), we were interested in analyzing potential influencing factors. Given that the procedure of the confidence forced-choice paradigm might rely on memory resources, we explored effects of short-term memory capacity, as assessed with the backward digit span measure, on meta-perceptual sensitivity. We found no evidence that individual differences in short-term memory capacity were linked to the average CMI across all conditions, *r*(54) = −.04, *p* = .791, the average unimodal CMI, *r*(54) = −.05, *p* = .740, nor the average crossmodal CMI, *r*(54) < .01, *p* = .975. Next, we scrutinized the idea that meta-perceptual ability is associated with cognitive control resources and that crossmodal comparisons might draw on more cognitive control resources. We did not find an association between individual executive abilities (as combined in an EF score for each participant) and average CMI across all conditions, *r*(54) = −.04, *p* = .791. When analyzed separately for each comparison type, neither average unimodal CMIs, *r*(54) = −.06, *p* = .669, nor crossmodal CMIs were linked to executive abilities, *r*(54) = .04, *p* = .788. Finally, we tested separately for each condition whether the difference in RT between the chosen and unsorted confidence set is associated with meta-perceptual sensitivity. For the unimodal comparison types, we found a marginally significant correlation with CMIs in the visual task, *r*(54) = .27, *p* = .050, and a significant correlation in the tactile task, *r*(54) = .32, *p* = .019. Interestingly, these correlations were absent for the crossmodal comparison type in both modalities, visual: *r*(54) = −.13, *p* = .338; tactile: *r*(54) = .11, *p* = .425. Thus, individual differences in processing dynamics cannot account for the observed high variability in crossmodal CMIs. But it appears that participants profit from RT differences for unimodal comparisons.

## Discussion

The present study addressed the question whether visual and tactile confidence share a common scale. Recent evidence already suggested that visual and tactile confidence underly a common mechanism (Faivre et al., 2018), but so far, this idea was only directly tested for vision and audition (deGardelle et al., 2016). Our findings indicate that confidence for a visual and tactile task is stored in an abstract, modality-independent format (Rouault et al., 2018) and is thus in support of confidence as a common currency. We found that participants were able to evaluate the quality of their perceptual decisions appropriately, even when the decisions were made across different sensory modalities. Importantly, confidence comparisons across the two modalities were as precise as comparisons within the same modality. This finding extends previous research showing that observers can compare their confidence across a visual and an auditive task without any additional costs (deGardelle & Mamassian, 2016). However, we also observed a slight bias towards the tactile modality (cf. Fairhurst et al., 2018): In the crossmodal conditions, some participants tended to select more trials from the tactile task as relatively more confident in comparison to the visual task. Though overall, this tendency to favor touch over vision in some cases did not hamper participant’s ability to properly compare their confidence across both modalities.

Furthermore, response times for the first-order perceptual decisions were slightly increased in the crossmodal blocks in comparison to the unimodal blocks. This increase was more pronounced for decisions in the visual task. It is likely that lengthened response times in the crossmodal blocks reflect task-switching costs (Kiesel et al., 2010) and it was also previously observed in the same paradigm but with a visual and auditory task (deGardelle et al., 2016). Response times for the perceptual decisions are used by some participants to form their confidence (Kiani et al., 2014) – longer response times could indicate longer processing and should thus be associated with lower confidence. We found that differences in response times were associated with meta-perceptual ability in the unimodal blocks but not in the crossmodal blocks. But although response time patterns were altered for crossmodal comparisons and participants could not use this information as efficiently as for unimodal comparisons, they were still able to use confidence as a common scale between the visual and tactile task.

## Acknowledgements

This work was funded by the German Research Foundation (Deutsche Forschungsgemeinschaft, DFG), Collaborative Research Centre SFB/TRR 135: Cardinal Mechanisms of Perception, project B5 and A4, project number 222641018. We thank Sabine Margolf and Maren Buck for help with data collection.

Commercial relationships: none

## References

Ais, J., Zylberberg, A., Barttfeld, P., & Sigman, M. (2016). Individual consistency in the accuracy and distribution of confidence judgments. Cognition, 146, 377–386. https://doi.org/10.1016/j.cognition.2015.10.006

Baer, C., & Odic, D. (2020). Children flexibly compare their confidence within and across perceptual domains. Developmental Psychology, 56(11), 2095–2101. https://doi.org/10.1037/dev0001100

Brainard, D. H. (1997). The psychophysics toolbox. Spatial Vision, 10(4), 433–436. https://doi.org/10.1163/156856897x00357

deGardelle, V., Le Corre, F., & Mamassian, P. (2016). Confidence as a common currency between vision and audition. PloS One, 11(1), e0147901. https://doi.org/10.1371/journal.pone.0147901

deGardelle, V., & Mamassian, P. (2014). Does confidence use a common currency across two visual tasks? Psychological Science, 25(6), 1286–1288. https://doi.org/10.1177/0956797614528956

Deroy, O., & Fairhurst, M. T. (2019). Spatial certainty : Feeling is the truth. In T. Cheng, O. Deroy, & C. Spence (Eds.), Routledge studies in contemporary philosophy: Vol. 122. Spatial senses: Philosophy of perception in an age of science (1st ed., pp. 183–198). Taylor & Francis, Taylor & Francis Group. https://doi.org/10.4324/9781315146935-11

Desender, K., Boldt, A., & Yeung, N. (2018). Subjective confidence predicts information seeking in decision making. Psychological Science, 29(5), 761–778. https://doi.org/10.1177/0956797617744771

Fairhurst, M. T., Travers, E., Hayward, V., & Deroy, O. (2018). Confidence is higher in touch than in vision in cases of perceptual ambiguity. Scientific Reports, 8(1), 15604. https://doi.org/10.1038/s41598-018-34052-z

Faivre, N., Filevich, E., Solovey, G., Kühn, S., & Blanke, O. (2018). Behavioral, modeling, and electrophysiological evidence for supramodality in human metacognition. The Journal of Neuroscience : The Official Journal of the Society for Neuroscience, 38(2), 263–277. https://doi.org/10.1523/JNEUROSCI.0322-17.2017

Fernandez-Duque, D., Baird, J. A., & Posner, M. I. (2000). Executive attention and metacognitive regulation. Consciousness and Cognition, 9(2), 288–307. https://doi.org/10.1006/ccog.2000.0447

Fleming, S. M., & Dolan, R. J. (2012). The neural basis of metacognitive ability. Philosophical Transactions of the Royal Society of London. Series B, Biological Sciences, 367(1594), 1338–1349. https://doi.org/10.1098/rstb.2011.0417

Fleming, S. M., Dolan, R. J., & Frith, C. D. (2012). Metacognition: Computation, biology and function. Philosophical Transactions of the Royal Society of London. Series B, Biological Sciences, 367(1594), 1280–1286. https://doi.org/10.1098/rstb.2012.0021

Fleming, S. M., & Lau, H. C. (2014). How to measure metacognition. Frontiers in Human Neuroscience, 8, 443. https://doi.org/10.3389/fnhum.2014.00443

Härting, C., Markowitsch, H. J., Neufeld, H., Calabrese, P., Deisinger, K., & Kessler, J. (2000). Deutsche Adaptation der revidierten Fassung der Wechsler-Memory Scale (WMS-R). Huber.

Kiani, R., Corthell, L., & Shadlen, M. N. (2014). Choice certainty is informed by both evidence and decision time. Neuron, 84(6), 1329–1342. https://doi.org/10.1016/j.neuron.2014.12.015

Kiesel, A., Steinhauser, M., Wendt, M., Falkenstein, M., Jost, K., Philipp, A. M., & Koch, I. (2010). Control and interference in task switching—a review. Psychological Bulletin, 136(5), 849–874. https://doi.org/10.1037/a0019842

Kleiner, M. (2010). Visual stimulus timing precision in psychtoolbox-3: Tests, pitfalls and solutions. Perception, 39, 189.

Koriat, A. (2007). Metacognition and consciousness. In P. D. Zelazo, M. Moscovitch, & E. Thompson (Eds.), Cambridge handbooks in psychology. The Cambridge handbook of consciousness (pp. 289–326). Cambridge University Press. https://doi.org/10.1017/CBO9780511816789.012

Kortte, K. B., Horner, M. D., & Windham, W. K. (2002). The trail making test, part b: Cognitive flexibility or ability to maintain set? Applied Neuropsychology, 9(2), 106–109. https://doi.org/10.1207/S15324826AN0902_5

Kreuzpointner, L., Lukesch, H., & Horn, W. (2013). Leistungsprüfsystem 2, LPS-2: Manual. Hogrefe.

Lau, H. C., & Passingham, R. E. (2006). Relative blindsight in normal observers and the neural correlate of visual consciousness. Proceedings of the National Academy of Sciences of the United States of America, 103(49), 18763–18768. https://doi.org/10.1073/pnas.0607716103

Mamassian, P. (2016). Visual confidence. Annual Review of Vision Science, 2, 459–481. https://doi.org/10.1146/annurev-vision-111815-114630

Mamassian, P. (2020). Confidence forced-choice and other metaperceptual tasks. Perception, 49(6), 616–635. https://doi.org/10.1177/0301006620928010

Mazancieux, A., Fleming, S. M., Souchay, C., & Moulin, C. J. A. (2020). Is there a g factor for metacognition? Correlations in retrospective metacognitive sensitivity across tasks. Journal of Experimental Psychology. General, 149(9), 1788–1799. https://doi.org/10.1037/xge0000746

Miyake, A., & Friedman, N. P. (2012). The nature and organization of individual differences in executive functions: Four general conclusions. Current Directions in Psychological Science, 21(1), 8–14. https://doi.org/10.1177/0963721411429458

Morales, J., Lau, H., & Fleming, S. M. (2018). Domain-general and domain-specific patterns of activity supporting metacognition in human prefrontal cortex. The Journal of Neuroscience : The Official Journal of the Society for Neuroscience, 38(14), 3534–3546. https://doi.org/10.1523/JNEUROSCI.2360-17.2018

Mueller, S. T., & Piper, B. J. (2014). The psychology experiment building language (pebl) and pebl test battery. Journal of Neuroscience Methods, 222, 250–259. https://doi.org/10.1016/j.jneumeth.2013.10.024

Palmer, E. C., David, A. S., & Fleming, S. M. (2014). Effects of age on metacognitive efficiency. Consciousness and Cognition, 28, 151–160. https://doi.org/10.1016/j.concog.2014.06.007

Pierce, C. S., & Jastrow, J. (1884). On small differences in sensation. Memoirs of the NationalAcademy of Science, 3, 75–83.

Reitan, R. M., & Wolfson, D. (1985). The Halstead–Reitan Neuropsycholgical Test Battery: Therapy and clinical interpretation. Neuropsychological Press.

Roebers, C. M. (2017). Executive function and metacognition: Towards a unifying framework of cognitive self-regulation. Developmental Review, 45, 31–51. https://doi.org/10.1016/j.dr.2017.04.001

Rouault, M., McWilliams, A., Allen, M. G., & Fleming, S. M. (2018). Human metacognition across domains: Insights from individual differences and neuroimaging. Personality Neuroscience, 1. https://doi.org/10.1017/pen.2018.16

Schütt, H. H., Harmeling, S., Macke, J. H., & Wichmann, F. A. (2016). Painfree and accurate bayesian estimation of psychometric functions for (potentially) overdispersed data. Vision Research, 122, 105–123. https://doi.org/10.1016/j.visres.2016.02.002

Stroop, J. R. (1935). Studies of interference in serial verbal reactions. Journal of Experimental Psychology, 18(6), 643–662.

Valk, S. L., Bernhardt, B. C., Böckler, A., Kanske, P., & Singer, T. (2016). Substrates of metacognition on perception and metacognition on higher-order cognition relate to different subsystems of the mentalizing network. Human Brain Mapping, 37(10), 3388–3399. https://doi.org/10.1002/hbm.23247

Wechsler, D. (2008). Wechsler adult intelligence scale – Fourth Edition (WAIS–IV). The Psychological Corporation.

